# Transcriptome data from silica-preserved leaf tissue reveals gene flow patterns in a Caribbean bromeliad

**DOI:** 10.1101/2023.06.16.545126

**Authors:** Natalia Ruiz-Vargas, Karolis Ramanauskas, Alexa S. Tyszka, Roberta J. Mason-Gamer, Joseph F. Walker

**Author notes:** = corresponding authors.

## Abstract

- Transcriptome sequencing is a cost-effective approach that allows researchers to study a broad range of questions. However, to preserve RNA for transcriptome sequencing, tissue is often kept under special conditions, such as immediate ultracold freezing. Here, we demonstrate that RNA can be obtained from six-month-old, field collected samples stored in silica gel at room temperature. Using these transcriptomes, we explore the evolutionary relationships of the genus *Pitcairnia* (Bromeliaceae) in the Dominican Republic and infer barriers to gene flow.
- We extracted RNA from silica-dried leaf tissue from 19 *Pitcairnia* individuals collected across the Dominican Republic. We used a series of macro-and micro-evolutionary approaches to examine the relationships and patterns of gene flow among individuals.
- We produced high-quality transcriptomes from silica-dried material and demonstrated that evolutionary relationships on the island match geography more closely than species delimitation methods. A population genetic examination indicates that a combination of ecological and geographic features are barriers to gene flow in *Pitcairnia*.
- High-quality transcriptomes can be obtained from silica-preserved tissue. The genetic diversity among *Pitcairnia* populations does not warrant classification as separate species, but the Dominican Republic contains several barriers to gene flow, notably the Cordillera Central mountain range.

## Introduction

The patterns of molecular evolution that shape the plant tree of life are rapidly coming to light owing, in large part, to advancements in and reduced cost of sequencing technologies. Whole genome sequencing often remains unaffordable for studies involving non-model organisms, but genome subsampling methods allow researchers to study these species (McKain et al., 2018). Among these methods are restriction site associated sequencing (Andrews et al., 2016; Eaton et al., 2017), target enrichment protocols (Johnson et al., 2019; Dodsworth et al., 2019), genome skimming (Zeng et al., 2018), and transcriptome sequencing (Wickett et al., 2014; Yang et al., 2017).

Transcriptome sequence data is easily transferable between studies and allows researchers to pursue a broad range of evolutionary questions, including inferring paleopolyploidy events (Jiao et al., 2011), uncovering putatively adaptive gene duplications (Brockington et al., 2015), and estimating population structure (Raduski et al., 2021). While most sequencing approaches for evolutionary studies of non-model organisms use DNA, transcriptome sequencing requires RNA, which is less stable and degrades faster. Therefore, transcriptome data is generally only obtainable from samples preserved in solutions such as RNA-later™ or flash-frozen in liquid nitrogen and maintained at temperatures of −70°C or lower (Logeman et al., 1987; Vennapusa et al., 2020). These sample preparation requirements have greatly limited the application of transcriptomic approaches, especially for taxa from regions where cryo-preservation resources are limited.

A recent study by He et al. (2022) found that the quality of transcriptomes obtained from plant tissue dried in silica gel and preserved at −20°C is comparable to liquid nitrogen frozen tissue. These results suggest that previously imposed limitations of tissue preservation may be overly cautious, and the study provided a framework for obtaining transcriptomes from a broader range of samples. To further explore this possibility, we investigated whether acceptable transcriptomes could be obtained from field collected plant tissue preserved in silica gel at room temperature for up to six months. These conditions replicate those of even the most prolonged fieldwork trips and test the feasibility of obtaining transcriptome data from any species in the world where collections are possible.

The focal group of this study are the flowering plants in the genus *Pitcairnia,* specifically those from the Dominican Republic. *Pitcairnia* is the second largest genus in the Neotropical flowering plant family Bromeliaceae, with over 300 species (Luther, 2014). A molecular phylogenetic analysis based on four chloroplast markers suggested that all Caribbean species of the genus are monophyletic (Schubert, 2017). Unlike other islands in the Caribbean, where each island contains a single species of *Pitcairnia*, Ayiti (Hispaniola^1^) contains up to six named species (Acevedo-Rodríguez & Strong, 2012), inferred from the high level of morphological disparity observed among individuals. *Pitcairnia* grows throughout the diverse landscape and varied niches of the Dominican Republic, which may have promoted speciation and led to the diversification and increased morphological disparity of *Pitcairnia*.

In this study, we use transcriptome data to examine the diversity of *Pitcairnia* native to the Dominican Republic. We demonstrate that transcriptome data for evolutionary studies is obtainable from field collected silica-dried tissue. In examinations of molecular diversity, we find that the morphology-based species delimitation in *Pitcairnia* of the Dominican Republic is not supported. We do uncover barriers to gene flow that correspond to the topography and ecological characteristics of the Dominican Republic, but they are not sufficient to lead to speciation. This study provides a framework for researchers seeking to obtain and analyze transcriptome data from field collected samples.

## Materials and Methods

### Data availability

Raw paired end sequence reads were deposited to NCBI Sequence Read Archive (SRA) archive, project id PRJNA982071. Analysis scripts can be found at https://github.com/karolisr/pitcairnia-dr-nrv. Assemblies, alignments, and trees are available at (https://zenodo.org/record/8021855).

### Tissue collection, RNA extraction, and sequencing

Leaf tissue was collected from 19 *Pitcairnia* individuals from across the distribution of the genus in the Dominican Republic (Figure 1). Samples were collected in 2021 from June 28–July 2 or September 11–14, with the corresponding permits from the Ministry of Environment of the Dominican Republic. Collections were immediately placed in resealable plastic bags filled with silica gel. After the collection trips, samples were stored at room temperature prior to RNA extraction.

**Figure 1.**
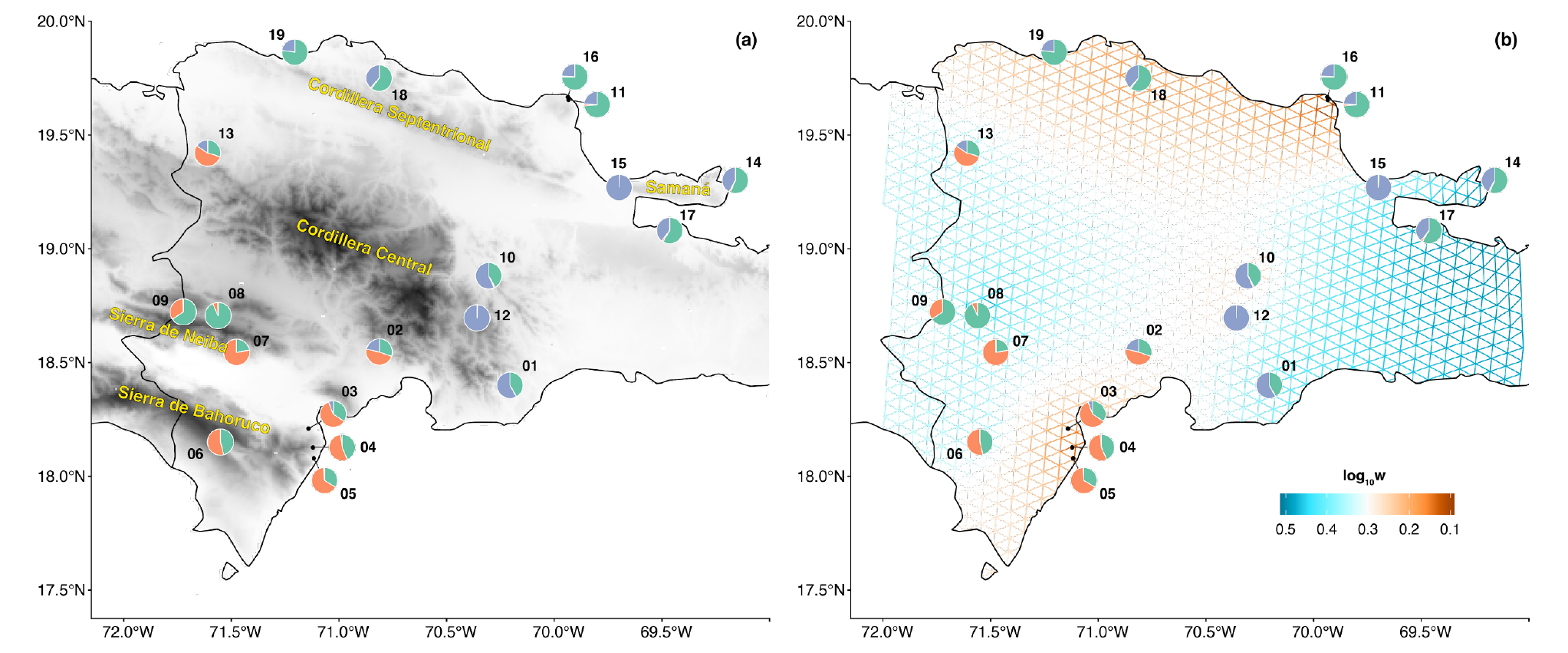
Maps of the Dominican Republic highlighting collections sites of *Pitcairnia* specimens. Pie charts are labelled with sample numbers and show the proportion of each layer inferred to have contributed to the ancestry of each sample based on the conStruct analysis. **(a)** Darker areas correspond to higher elevations, with major mountain ranges labeled. **(b)** The Fast Estimation of Effective Migration Surfaces (FEEMS) analysis. Warmer colors correspond to areas inferred to have less gene flow.

Either three or six months after collection, approximately 40 mg of tissue from each sample was ground to a fine powder in 1.5 mL tubes submerged in liquid nitrogen. RNA was extracted from the ground leaf tissue using the Sigma-Aldrich Spectrum™ Plant Total RNA Kit with minor modifications (Figure S1) and stored at −80°C. The University of Illinois at Chicago Sequencing Core assessed RNA quality using the RNA ScreenTape. Library preparation was performed using unstranded paired-end Total RNAseq with rRNA depletion. Paired-end 150 base-pair (bp) sequencing was carried out on a single lane of Illumina NovaSeq 6000 System, flow cell type S4, also at the UIC Sequencing Core.

Raw sequence reads for two outgroup species, *Pitcairnia albiflos* (SRR2518104) and *Pitcairnia staminae* (SRR2518089), were obtained from the NCBI SRA database using the fastq-dump utility from the SRA-Toolkit package version 3.0.0 (https://hpc.nih.gov/apps/sratoolkit.html).

### Morphological data

Specimens for the field collected samples were pressed and deposited in the National Herbarium of the Dominican Republic (JBSD) (Table S1). Trichome density, leaf width and length, and marginal spine size were measured because they were the most variable among collection sites. Furthermore, trichome density is considered important for specimen identification (Teodoro Clase, pers. comm. December 2022). Trichome density was estimated in the field and classified into three categories: High, Medium, and Low. Leaf width, length and marginal spine size were measured from the dried specimens. The average measurements of the three largest leaves is reported.

Two approaches were used to classify the samples: the first was to have a taxonomist specialized in Dominican flora assign species names to the collections. The second was to use the species names given to the populations based on geolocation data from herbarium specimens. Geolocation based assignment was unable to classify Samples 13 and 19 because other herbarium specimens from their location were identified only to genus (i.e., *Pitcairnia sp.*). Sample 13 also could not be classified by the taxonomist and consequently remained undetermined as *Pitcainia sp.* The geolocation based classification identified four samples of *Pitcairnia elizabethae*, four *P. fuertesii*, three *P. domingensis*, two *P. ariza-juliae*, two *P. samuelsonii*, and two *P. jimenezii*. In contrast, the taxonomist classification identified eight *P. elizabethae*, seven *P. dominingensis*, and three *P. fuertesii*.

### Read processing and transcriptome assembly

Random sequencing errors in the raw reads were identified and corrected using Rcorrector v1.0.4 (Song and Florea, 2015). Uncorrectable reads were removed using the unfixable_filter.py script from Morales-Briones et al. (2021). Trimmomatic v0.39 (Bolger et al., 2014) was used to trim and remove low-quality sequences with the settings “SLIDINGWINDOW:4:5 LEADING:5 TRAILING:5 MINLEN:25”. Kraken2 v2.1.2 (Wood et al., 2019) was used to filter human, bacterial, protozoan, and fungal reads using the Kraken2 PlusPF database (https://genome-idx.s3.amazonaws.com/kraken/k2_pluspf_20220607.tar.gz) with default settings. Kraken2, with a confidence value of 0.1, was further used to remove plastid and mitochondrial reads using their respective pre-made databases from Kakapo (Ramanauskas and Igic, 2023). All reads not identified as contamination or organelle genomes were retained as nuclear reads.

For each sample, the nuclear reads were assembled using Trinity v2.11 (Haas et al., 2013) with default settings and the option “--no_normalize_reads”. Chimeric sequences were removed following the Yang and Smith (2013) procedure, using Blastx v2.13.0 (Altschul et al., 1990) with a protein database composed of the *Arabidopsis thaliana* (TAIR 10), *Oryza sativa* (IRGSP 1.0), *Zea mays* (B73 REFERENCE-NAM-5.0), and *Ananas comosus* (F153) coding sequences, hereafter AOZA database, downloaded from PlantEnsembl (Bolser et al., 2016). The program Corset v1.09 (Davidson and Oshlack, 2014), with mapping conducted using Salmon v0.9.1 (Patro et al., 2017), was used to filter for the best transcript. The open reading frames were predicted using Transdecoder v5.5.0 (https://github.com/TransDecoder/TransDecoder) and predictions were guided by the AOZA database. To further remove possible contamination, only genes with 70% nucleotide sequence similarity, as inferred by Blastn v2.13.0, compared to the corresponding nucleotide sequences in the AOZA database, were retained.

The same procedure was conducted for both the plastid and the mitochondrion datasets to infer organelle transcriptomes, with a database composed of (NC_026220.1: *Ananas comosus*, NC_000932.1: *Arabidopsis thaliana*, NC_031333.1: *Oryza sativa*, and NC_001666.2: *Zea mays*) for plastid and (NC_007982.1: *Zea mays*, NC_007886.1: *Oryza sativa*, and NC_037304.1: *Arabidopsis thaliana*) for the mitochondrion. For each predicted organelle transcriptome, the longest single transcript corresponding to an organelle protein coding sequence was retained for downstream analysis.

### Organelle phylogenetic analyses

The nucleotide sequences for individual plastid protein coding sequences were aligned using Prank v170427 (Loytynoja and Goldman, 2008) with default settings. Alignments were cleaned for a minimum column occupancy of 10% using the Phyx v1.2 (Brown et al., 2017) program *pxclsq*. All genes were then concatenated using the Phyx program *pxcat*, and the plastid tree was inferred using maximum likelihood as implemented in IQTREE v1.6.12 (Nguyen et al., 2015) with default model selection, the proportional partition model (Chernomor et al., 2016) and 1000 UFBoot2 (Hoang et al., 2018) replicates. The same procedure was performed for the mitochondrion protein coding sequences.

### Nuclear phylogenetic analyses

Cd-hit v4.8.1 (Fu et al., 2012), with the settings “-c 0.99 −n 10 −r 0”, was used to reduce redundancy in the nuclear transcriptomes. All-by-all Blastn was used to identify sequence similarity of combined transcriptomes setting the e-value to 10 and max-target-seqs to 1000. The clusters of similar sequences were refined using mcl v14-137 (Van Dongen, 2000) with an inflation value of 1.4. Only clusters with at least four taxa, hereafter referred to as homolog clusters, were retained.

Mafft v7.490 (Katoh and Standley, 2013) with the settings “--auto --maxiterate 1000” was used to individually align the homolog clusters. The resulting alignments were cleaned with *pxclsq* to ensure a minimum column occupancy of 10%. Homolog trees were inferred using IQ-TREE with 1000 UFBoot2 replicates and automated model selection. Long branches with relative values of 0.02 substitutions per site (subs/site) or more, that were 10x longer than the sister branch, and/or had an absolute value of 0.03 subs/site or more were removed from the homolog tree. Any homolog tree with a branch longer than 0.2 subs/site and at least four taxa on both sides of the branch was split on the branch and treated as separate homologs for downstream analysis. The same process with the same settings was repeated a second time.

Orthology was inferred using the Maximum Inclusion method (Yang and Smith, 2014) setting the relative value to 0.02 subs/site, the absolute value to 0.03 subs/site and requiring a minimum of 10 taxa required for each ortholog. Ortholog alignments were extracted from the alignments used to infer homolog trees in the same manner as Wheeler et al. (2022). Individual gene trees were inferred using IQ-TREE with default model selection and 1000 UFBoot2 replicates. The individual relationships were estimated using the maximum quartet support species tree method as implemented in Astral v5.7.8 (Zhang et al., 2018). Support values for the Astral tree were inferred using the coalescent-based quartet frequency method (Sayyari and Mirarab, 2016). Molecular branch lengths were applied to the Astral tree using the quadripartition concordance approach as implemented in Bes (Walker et al., 2022).

### Variant-calling workflow

The outgroup taxa (*P. albiflos* and *P. staminae*) were included in the variant calling procedure. A reference was selected for each ortholog by identifying the longest sequence from among the ingroup taxa. STAR v2.7.10a (Dobin et al., 2013) was used to map the processed sequence reads from each sample to the reference with the setting “--twopassMode”. The Picard command line tools (Broad Institute, 2018) were used to filter potential PCR duplicate reads from each alignment. The genotypes, including single nucleotide polymorphisms (SNPs) and invariant sites, were identified using FreeBayes v1.3.6 with all samples assigned to a single population (Garrison and Marth, 2012). The genotype calls were filtered by removing sites that met any of the following criteria: (1) the site was inferred to have more than two alleles segregating across all samples, (2) the site was called polymorphic but had a “Phred-scaled quality score” (QUAL field in the VCF file; 10log10prob[no variant]) lower than 20, (3) the sites read depth was lower than 10 in any of the 19 samples, (4) the site had one allele overrepresented in heterozygous samples (allele balance; AB < 0.25 or AB > 0.75), (5) the site had AB > 0.05 in homozygous samples, or (6) the site had variants with a frequency lower than 0.05 across all samples (minor allele frequency, MAF ≤ 0.05).

### Principal component analysis

The R package SNPRelate v1.32.1 (Zheng et al., 2012) was used to filter genotypes and perform a principal component analysis (PCA) (McVean et al., 2009). Linkage disequilibrium (LD) was filtered from the genotypes using the function snpgdsLDpruning with LD threshold set to zero (i.e. one SNP per locus was used) and the missing sample rate threshold set to one sample (ld.threshold=0.0, missing.rate=1/19 + 1e-10). The PCA analysis was conducted using the function snpgdsPCA.

### Genetic structure analysis using conStruct

Genetic structure analysis was performed using R package conStruct v1.0.5 (Bradburd et al. 2018). Only loci present in at least 15 of the 19 samples were used. Potential linkage disequilibrium bias was reduced by randomly choosing one SNP per locus. Samples were not assigned to populations, *i.e.* each sample was treated as a separate population. After running cross-validation, a spatial model with three layers (k=3) was chosen and a run with four chains of 20,000 generations was performed. Spatial information for the conStruct analysis was incorporated using the geographic coordinates of the collected samples.

### Fast Estimation of Effective Migration Surfaces (FEEMS)

A Fast Estimation of Effective Migration Surfaces (FEEMS) (Marcus et al., 2021) analysis was performed to infer and visualize potential barriers to gene flow. The same set of SNPs was used for this analysis as for the conStruct analysis. The optimal smoothness parameter (*λ* value) was inferred using cross-validation. An outline of the island of Ayiti was created manually using the online *Polyline* tool (https://www.keene.edu/campus/maps/tool). A Geodesic Discrete Global Grid Systems grid intersecting the island’s outline was generated using the R package dggridR (Barnes and Sahr, 2017). Grid size must be selected by a user and, therefore, it is somewhat arbitrary. FEEMS assigns samples to a closest node on the grid. If a grid is too large all samples will be assigned to a single node. On the other hand, performance increases nonlinearly with increasing node density. When considering these factors we chose a grid size which assigned any samples with a pairwise distance above 5 km to a unique node.

## Results

### RNA extraction quality assessment and transcriptome assembly

Across the 19 samples, the RNA integrity (RIN) values ranged from 2.3 to 7.4 (Table 1; Table S2). The number of sequenced reads ranged from a minimum of 77,268,317 for Sample 02, to a maximum of 99,484,187 for Sample 01. Slightly less than half of the reads from each sample were filtered as either human, protozoan, bacterial or fungal contamination. Of the remaining reads for Sample 01: 46,116,739 reads were predicted to be nuclear, 865,000 were predicted to be plastid, and 1,407,674 were predicted to be mitochondrion. For Sample 02: 46,578,119 reads were predicted to be nuclear, 6,349,726 reads were predicted to be plastid, and 1,682,695 reads were predicted to be mitochondrion. After a final filtering step that required all predicted transcripts to have at least 70% nucleotide similarity with the coding sequences of *Arabidopsis thaliana*, *Oryza sativa*, *Zea mays*, or *Ananas comosus*, we found that Sample 01 had a predicted 23,185 nuclear genes, 37 chloroplast genes, and 36 mitochondrial genes, and Sample 02 had 24,990 nuclear genes, 38 chloroplast genes, and 39 mitochondrial genes.

**Table 1.** Summary results of the RNA extractions and assembled transcriptomes used in the study. *Pitcairnia albiflos* and *P. staminae* were downloaded from the NCBI short read archive and therefore do not Total RNA or RIN values.

| Sample Name | Total RNA (ng) | RIN | Total sequence reads | Nuclear sequences | Chloroplast sequences | Mitochondrial sequences |
| --- | --- | --- | --- | --- | --- | --- |
| Sample 01 | 510 | 2.3 | 99,484,187 | 46,578,119 | 6,349,726 | 1,682,695 |
| Sample 02 | 840 | 4.3 | 77,268,317 | 46,116,739 | 865,000 | 1,407,674 |
| Sample 03 | 528 | 3.8 | 81,670,840 | 45,084,923 | 927,284 | 1,320,711 |
| Sample 04 | 4930 | 5.4 | 77,788,478 | 51,905,175 | 3,207,685 | 2,675,367 |
| Sample 05 | 510 | 3.2 | 89,263,036 | 49,957,936 | 5,133,436 | 1,909,637 |
| Sample 06 | 390 | 4.3 | 99,040,548 | 65,979,048 | 2,596,433 | 3,218,196 |
| Sample 07 | 2380 | 3.7 | 87,455,923 | 46,613,054 | 8,306,422 | 2,209,659 |
| Sample 08 | 1581 | 7.4 | 95,356,788 | 56,232,278 | 99,537 | 2,066,630 |
| Sample 09 | 6600 | 4.8 | 86,434,374 | 49,579,100 | 3,901,460 | 1,678,802 |
| Sample 10 | 3100 | 4.1 | 80,217,537 | 49,375,135 | 3,585,535 | 2,127,142 |
| Sample 11 | 1020 | 5.7 | 90,185,382 | 45,720,319 | 866,775 | 1,421,561 |
| Sample 12 | 600 | 5 | 86,422,214 | 60,871,492 | 2,105,689 | 1,678,563 |
| Sample 13 | 1110 | 5 | 98,358,395 | 58,213,213 | 3,565,910 | 2,246,538 |
| Sample 14 | 899 | 4.9 | 95,451,297 | 52,588,180 | 1,376,271 | 1,884,013 |
| Sample 15 | 690 | 3.7 | 81,388,060 | 50,583,456 | 4,039,181 | 2,913,524 |
| Sample 16 | 990 | 4.1 | 84,733,604 | 49,143,522 | 4,013,399 | 2,781,999 |
| Sample 17 | 1488 | 5.7 | 85,161,150 | 45,201,696 | 478,966 | 1,656,652 |
| Sample 18 | 406 | 4.2 | 79,558,466 | 45,963,594 | 974,380 | 2,267,451 |
| Sample 19 | 1290 | 6.1 | 88,947,285 | 52,751,861 | 1,804,642 | 1,206,667 |
| *P. albiflos* | NA | NA | 56,976,403 | 40,661,866 | 325,139 | 46,110 |
| *P. stamineae* | NA | NA | 51,790,009 | 43,995,454 | 57,717 | 9,067 |

### Inferred relationships among individuals

We recovered 65 different plastid protein-coding sequences from the transcriptomes, 53 of which were present in at least four samples (Table S3). The transcriptome data provided 63 different mitochondrion protein coding sequences (genes or open reading frames), 55 of which were present in at least four samples (Table S4).

The majority of relationships were poorly supported (UFboot <= 95%) for both organelle phylogenies. The individuals did not form clades corresponding to “species” based on taxonomic determinations (Figure S2 and S3). Most relationships in the mitochondrion tree conflicted with relationships in the plastid tree (Figure S2 and S3). In the mitochondrion tree, the outgroups, *Pitcairnia staminae* and *Pitcairnia albiflos,* were not monophyletic. The mitochondrion tree length (i.e., the sum of all branch lengths) was 0.286 subs/site compared to the plastid tree length of 0.751 subs/site. The total parsimony informative characters for the plastid dataset was 2,698 sites compared to 1,060 sites for the mitochondrion dataset (Figure S4).

We recovered 11,683 nuclear orthologs shared by at least 10 of the 21 samples (19 ingroups and 2 outgroups). The relationships in the nuclear tree did not correspond to species identifications from either the taxonomist’s determinations or species assignments based on geolocation data, nor were they concordant with either organelle tree. However, unlike the organelle trees, the nuclear relationships were well supported, with nearly all relationships containing local posterior probabilities (>= 0.95). The structure of the nuclear phylogeny was largely concordant with the relationships expected based on the island’s ecology and geography (Figure 2).

**FIGURE 2.**
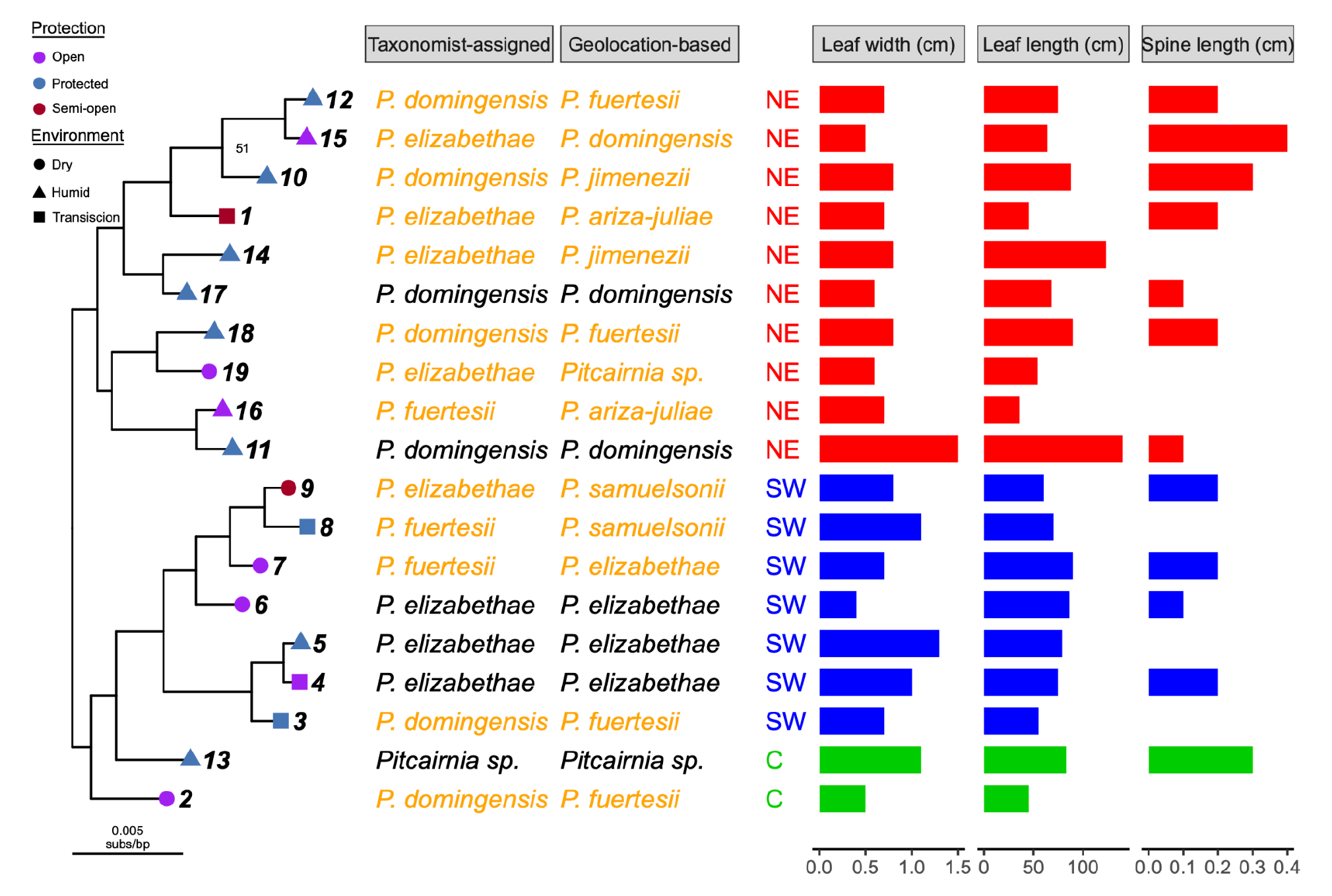
Phylogenetic relationships inferred from 11,683 nuclear gene trees, taxonomic classifications and morphological traits for 19 samples of *Pitcairnia*. The phylogenetic relationships among individuals do not correspond to taxonomic classifications or morphological traits. The tree was inferred with a coalescent-based maximum quartet species tree method and support for the relationships are local posterior probabilities (LPP). The LPP values for all poorly supported relationships (LPP < 95%) are labeled. Branch lengths are in units of substitutions per site and the values depicted are the mean lengths of corresponding branches from concordant gene trees. The shapes and colors on the tips of the tree correspond to the environmental conditions of the sample location and the number corresponds to the individual sample number. The species names were assigned in two ways: by a taxonomic specialist and based on the geolocation data. Assigned species names are aligned with the corresponding tree tip labels, as are three major morphological characters used to help guide taxonomic identification. The colors of the bar charts correspond to each sample’s region of origin.

### Variant Calling and Principal Components Analysis

FreeBayes called 492,539 Single Nucleotide Polymorphisms (SNPs) across the 11,683 nuclear loci (orthologs used in the phylotranscriptomic analysis). After the data were filtered using the criteria outlined in the materials and methods, 10,218 high-quality SNPs across 2,933 loci remained. The fraction of heterozygotes across the alleles followed a distribution similar to what would be expected under Hardy-Weinberg Equilibrium (Figure S5). To account for linkage disequilibrium and avoid SNP dense loci being more influential, we randomly retained one SNP per locus and ensured that the SNP was informative (i.e., variable across samples). The final result was 1,900 informative SNPs after all filtering steps.

In our principal component analysis, PC1 and PC2 explained 13.98% and 10.38% of the variation, respectively (Figure 3). The samples did not form tight clusters in the PCA; however, the samples collected from the North East (NE) side of the island aligned on the PC1 axis but not the PC2 axis, and vice versa for the South West (SW) side. Samples 13 and 02, collected from the island’s center, fell in the center of the graph.

**FIGURE 3.**
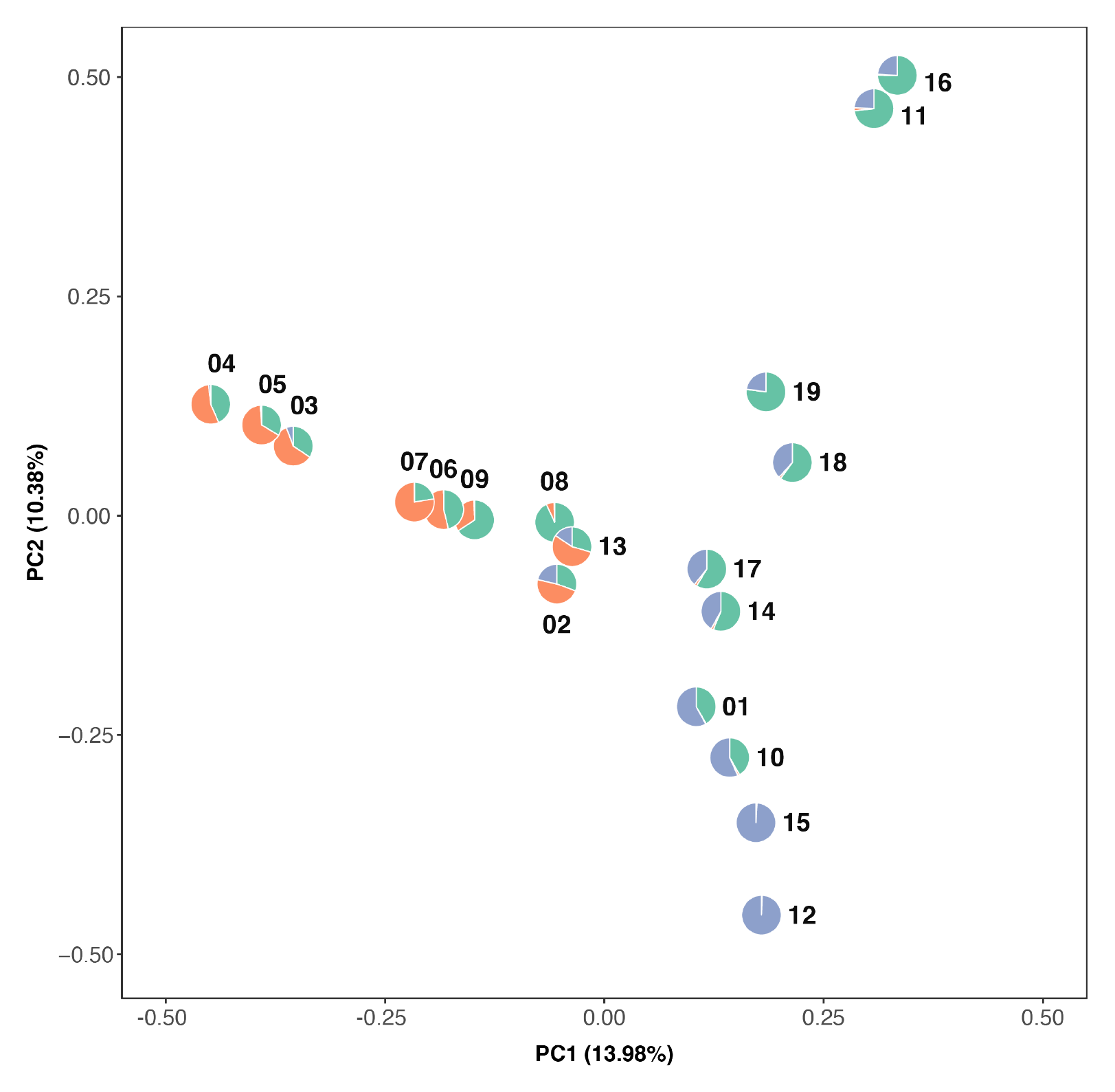
Principal component analysis of *Pitcairnia* individuals. The sample numbers for each individual are shown and the pie charts correspond to the proportion of each layer inferred to have contributed to the ancestry of the sample based on the conStruct analysis (as in Figure 1). Samples from the Northeast and Southwest sides of the Cordillera Central were differentiated by the first principal component.

### Analysis of genetic diversity across the Dominican Republic

The Fast Estimation of Effective Migration Surfaces (FEEMS) visualization showed a barrier to gene flow between the samples found on the NE and the SW side of the Cordillera Central mountain (Figure 4). The NE/SW barrier was corroborated by the conStruct analysis, which predicted a unique layer on each side of the mountain (Figure 1). Samples 02 and 13 were collected between the NE/SW sides and were inferred to have a mixture of the NE and SW layers based on the conStruct analysis, with a greater proportion from the SW layer.

Samples 03, 04, and 05, collected on the eastern side of the Sierra de Bahoruco mountain range, are the geographically and genetically closest individuals on the SW side of the Cordillera Central and show decreased gene flow with Sample 06, collected from the opposite side of the Sierra de Bahoruco. The FEEMS results do not suggest any decrease in gene flow by distance between Sample 13 and Samples 06, 07, 08 and 09. It does show decreased gene flow by distance from Sample 02 to the other SW samples.

On the NE side, FEEMS shows gene flow between samples 01, 10 and 12 in the Cordillera Central and Samples 14, 15, and 17 in the Samaná Peninsula and Bay, but decreased gene flow between those and Samples 11, 16, 18, and 19. Gene flow also appears to be reduced between the cluster of 11 and 16 and the cluster of 18 and 19, even though the individuals in both clusters show higher genetic similarities between each other than with any other samples.

## Discussion

### High quality transcriptomes may be obtained from silica-preserved tissue

This study demonstrates that RNA can still be successfully extracted and sequenced if the plant samples are stored in silica-gel at room temperature. The plant tissue was never frozen after collection, differentiating this study from previous work extracting RNA from silica-dried samples (He et al., 2022), which showed that storing silica-dried plant tissue at −20°C is sufficient for high-quality RNA extractions. The RIN values we obtained were generally lower than those recovered by He et al. 2022, and Sample 01’s RIN value of 2.3 is considered low quality. However, the assembled transcriptome for Sample 01 was of similar quality to those of samples with superior RIN values (Table 1; Table S2).

Historically, the stability of RNA prior to extraction has been a concern for transcriptome sequencing. However, we found that RNA in dried leaf material remains stable enough for short-read sequencing for at least six months, suggesting that transcriptomes may be obtained from plant tissues anywhere field collection is possible. The misconceptions about RNA stability may arise from differential gene expression studies, where there is a need to quickly stop cellular processes, or from the desire to retain long sequences for PCR (Natarajan et al., 2000). These misconceptions are likely reinforced by protocols requiring that tissue is frozen or stored in an RNA preservation solution (Logeman et al., 1987; Yang et al., 2017; Vennapusa et al., 2020). Future work has the potential to provide valuable insight into the length of time RNA remains intact and capable of providing transcriptomes, and to uncover any phylogenetic patterns in the rate of RNA degradation.

### Morphology and collection location provide conflicting species delimitation for *Pitcairnia*

There are six named species of *Pitcairnia* endemic to the Dominican Republic (DR). Of these, five species have been formally described (Acevedo-Rodríguez & Strong, 2012): *P. domingensis*, *P. elizabethae*, *P. fuertesii*, *P. jimenezii*, and *P. samuelsonii.* The sixth, *P. ariza-juliae*, was identified as new from herbarium specimens by Robert W. Read from the Smithsonian Institution (JBSD: Liogier 27724; Zanoni 15005; Zanoni 22952) but never formally described. Although this is not a published name, the specimens listed above are identified as such in the herbarium specimens. The species in Ayiti were described by Carl Mez (*P. fuertesii*) and Lyman B. Smith (all others) between 1913 and 1964 from pressed specimens (Smith & Downs, 1974).

The Flora Neotropica monograph of the Pitcairnioideae subfamily (Smith & Downs, 1974) is still the most comprehensive work done on the group. However, the monograph does not address how the taxa are different from each other, and the identification key (which covers all species of *Pitcairnia*) does not group the five described species together in the same subkeys. *Pitcairnia elizabethae* is characterized as the shortest, reaching a maximum height of 28 cm, and *P. fuertesii* and *P. jimenezii* as the tallest, both reaching heights of 100 cm. *Pitcairnia domingensis* and *P. jimenezii* are glabrous, while the rest are densely covered in trichomes on the abaxial side of the leaves. *Pitcairnia jimenezii* has no marginal spines while the rest have spines of varying length and density. The monograph does not describe any diagnostic characters that are unique to individual species. From our field observations, the most pronounced morphological disparities are in the length and width of leaves, the length of the flower stalk, the density of trichomes covering the leaves, and the density and size of marginal spines (Figure S6 and Figure S7).

The distribution of each species was thoroughly documented and outlined by Smith & Downs (1974). *Pitcairnia fuertesii* is the only species widely distributed across Ayiti, with specimens from the Samaná Peninsula, Sierra de Bahoruco and the Cordillera Central mountain ranges in the DR and Massif du Nord and Massif de la Selle mountain ranges in Haiti. *Pitcairnia domingensis* is circumscribed to Samaná, *P. elizabethae* from Massif du Nord in Haiti and Sierra de Bahoruco in the DR, *P. jimenezii* from Puerto Plata Province, and *P. samuelsonii* from the Massif du Nord and Massif de la Selle in Haiti and the San Juan de la Maguana province in the DR. Since then, new records for each species have been recorded in museum specimens.

The lack of an identification key specific to the species from Ayiti or the Caribbean have made the identification of *Pitcairnia* specimens difficult. To account for this and maximize the diversity of collections, sampling in this study covered the genus’s historically reported range across the DR. Some locations in the center of the Cordillera Central have been altered by human development and *Pitcairnia* was not found during field work.

### Phylogenetic analyses of the nuclear and organellar genomes do not support species delimitations

The two methods of species identification (taxonomist and geolocation) disagreed for twelve of the seventeen samples, highlighting the need to examine the relationships further. None of the measurements performed on the samples (trichome density, size of marginal spines, and leaf length and width) corroborate either classification scheme (Figure S6). As *Pitcairnia* species are delimited based on quantitative traits, it is possible that these differences are the result of plasticity, which may be exacerbated by the diverse ecology of the DR. Molecular phylogenies can often help with species delimitation, as molecular data provides a large number of characters with which to estimate relationships.

The plastome and mitochondrial genomes remain powerful tools for molecular phylogenetics; they are two of the most character-rich markers, are often considered to each have a single evolutionary history, and are obtainable from transcriptome data (Morales-Briones et al., 2021). Based on a four plastome gene molecular phylogeny, the *Pitcairnia* of the Dominican Republic were inferred to be monophyletic (Schubert, 2017). In our dataset, each organelle contained more than 50 phylogenetically informative protein coding sequences. However, based on the low support for relationships, the organellar genomes proved insufficient for reliably estimating species relationships. Mitochondrial and plastid trees often have concordant topologies (Tyszka et al., 2023), but in our case, the lack of resolution and conflict between the two provide additional evidence that the relationships inferred by the organelle trees should not be trusted. Further examination of the amount of molecular evolution between samples shows that the topologies are inferred from limited information, as reflected by the small total tree length and low level of parsimony informative characters.

We also examined nuclear genome-based relationships using transcriptome data. One of the difficulties for inferring relationships using nuclear data is phylogenetic conflict, specifically Incomplete Lineage Sorting (ILS). Because *Pitcairnia* appears to have experienced a rapid radiation on the island Ayiti one might expect there to be high levels of ILS. To accommodate ILS, we used the coalescent-based maximum quartet support species tree method. This approach resulted in a species tree where almost all relationships were well-supported (Figure 2). Despite the strong support, the individuals did not form clades with their respective species, so neither classification method was validated. Instead of clustering based on taxonomic species determinations, the individuals clustered based on geography and ecology. Specifically, a major split in the tree was found between samples on the NE and the SW side of the Cordillera Central.

### Mountain ranges in the Dominican Republic are barriers to gene flow among *Pitcairnia* individuals

Based upon the short molecular branch lengths of the inferred phylogenies, indicating minimal molecular evolution among individuals, the data appeared suitable for population genetic analyses. Previous work has shown that transcriptomes can provide a data-rich source to infer patterns of gene flow (Raduski et al., 2021). Therefore, we proceeded with a series of population genetic analyses to further investigate the relationships among the individuals. In the principal component analysis (PCAs), the individuals did not form clear clusters with respect to the first two PC axes (Figure 3), indicating that there were not clearly differentiated groups that could be binned into separate populations for downstream analyses.

The program conStruct (Bradburd et al., 2018) was used to analyze gene flow across the island. ConStruct assumes several distinct ancestral populations called layers. Each individual is inferred to have some proportion of ancestry from each layer and the more similar the individuals, the more similar the composition of layers will be. Unlike other approaches to analyzing population structure, conStruct incorporates spatial information into the analysis. A complementary approach to analyzing and visualizing isolation by distance was conducted using Fast Estimation of Effective Migration Surfaces (FEEMS). FEEMS complements conStruct as it is robust to sparse sampling and functions well under simulated coalescent processes (Marcus et al., 2021). FEEMS determines whether the amount of genetic distance between individuals is what would be expected based on geographic distance. Any deviations from that may indicate a barrier to gene flow.

The split in the phylogeny corresponding to whether samples were collected NE or SW of the Cordillera Central was also reflected in the population genetic analyses (the PCA, conStruct, and FEEMS). The Cordillera Central is the tallest mountain range in the Caribbean and is a well-documented barrier to gene flow in some palms (Rodríguez-Peña *et al*., 2014) and a predicted driver of speciation in *Podocarpus* (Nieto-Blázquez et al., 2021). Based on all analyses, the Cordillera Central appears to be the best explanation for the genetic differentiation between the SW and the NE *Pitcairnia* individuals. All analyses performed resulted in concordant results, indicating robust support for this result regardless of the approach used.

On the NE side of the Cordillera Central, the relationship between species largely matches geographic distances; for example, Samples 18 and 19, and Samples 11 and 16 cluster together, both in the PCA and phylogeny. A comparison of Samples 11 and 16, in particular, illustrates how phenotypic plasticity might be responsible for the morphological differences observed between individuals (Figure S8). These two samples are the least genetically differentiated of any pair of samples and were collected 1.31 linear km apart, but they show greater differences in the quantitative morphological traits. The leaves of sample 11 reach 140 cm, were softer, and appeared greener in color because of a reduced density in trichomes. When these plants were collected, they were growing on the side of the road, and although close to and facing the ocean, they were found under lush vegetation. Sample 16 was shorter, with coarse, grayish leaves reaching 35 cm, and was growing on the cliffs directly facing the ocean.

For samples from the island’s SW side, a similar pattern of isolation by distance was observed. However, despite Sample 06 being geographically closer to Samples 03, 04, and 05, it was genetically closer to Samples 07, 08, and 09. The shared ecology may explain the genetic similarity, as Samples 06, 07, 08, and 09 were from drier habitats on the southern side of the Sierra Neiba mountain range, as compared to the wetter habitats of 03, 04 and 05. A pattern of decreased gene flow due to the Sierra de Neiba and Sierra de Bahoruco has been previously documented in the small mammal *Solenodon* (Turvey *et al*., 2016) and the cactus *Leptocereus* (Majure *et al*., 2021).

## Conclusions

This study provides several advances for transcriptomics in evolutionary biology. We found that RNA for short-read sequencing may be extracted from leaf samples dried in silica gel and stored at room temperature for up to six months. Using this transcriptome sequence data, we demonstrate that the *Pitcairnia* of the Dominican Republic does not show the predicted genetic isolation expected for multiple species, and the Cordillera Central is the main isolator to gene flow. These results demonstrate promise for obtaining transcriptomes from any plant growing where tissue collection is possible.

## Author Contributions

All authors helped design the experiment. NRV performed fieldwork and sample collection. NRV, KR, AST, and JFW performed the RNA extractions. Data analysis was performed by KR, NRV, and JFW, with assistance from AST. KR, NRV, and AST generated the figures for the manuscript with input from JFW and RJMG. NRV and JFW led the writing of the manuscript, with input from KR, AST, and RJMG. All authors approved the final draft of the manuscript.

## Acknowledgements

The authors would like to thank Ernesto Payano and the personnel of the Botanical Garden of Santo Domingo for assistance during field and herbarium work, in particular Pedro Toribio, and Teodoro Clase for specimen identification. May Yeo and Edwige Moyroud for sharing their adjustments to the RNA extraction protocol. Andy Randuski and Silas Tittes for advice and assistance on the data analysis. Gideon Bradburg for advice on ConStruct. Robyn Phillips, Drew Larson, Weston Testo, and Rosa Rodríguez-Peña for constructive criticism and helpful feedback on the manuscript. This work was supported by the Field Museum’s Negaunee Fieldwork Fund, startup funds from the University of Illinois, Chicago to JFW and National Science Foundation awards GRFP 2236870 to AST.

## Footnotes

1 Ayiti is the name given to the island by the Taino people, the indigenous group present at the time of colonization. The spelling differentiates it from the country of Haiti, located on the island.

