## SupplementaryMaterials for "Transcriptome data from silica-preserved leaf tissue reveals gene flow patterns in a Caribbean bromeliad"

### Supplementary Material

**Figure S1. RNA extraction protocol using Sigma Spectrum™ Plant Total RNA Kit.** Listed below are the changes made and the respective steps they correspond to from the Sigma Spectrum™ procedure. Aside from the listed changes, the protocol is the same as the Sigma Spectrum™ Plant Total RNA Kit procedure performed using protocol A with the On Column DNase digest.

**Grinding tissue** - Transfer approximately 40 mg of silica-preserved tissue into a 2 mL tube submerged in liquid nitrogen. Allow the tissue to sit in the 2mL tube for around 5 seconds, and then grind the tissue with a pestle.

**Pellet cellular debris** – centrifuge the samples at maximum speed for 5 minutes.

**Filter lysate** – centrifuge at maximum speed for 2 minutes.

**First column wash (WS1)** – add 300ul of wash solution 1 into the column.

**Second column wash (WS1)** - Centrifuge at maximum speed for 1 minute.

<u>Perform 4 column washes instead of three. The fourth column wash procedure is the same as the third column wash.</u>

**First elution –** Pipette 40ul nuclease free water.

<u>Perform the second elution</u>

**Figure S2. Inferred relationships among individuals using plastome data.** The tree was inferred using an ML concatenated supermatrix, and support for the relationships is based on Ultrafast bootstraps, with the UFboot values for all poorly supported relationships (UFboot < 95%) labeled. The shapes and colors on the tips correspond to the environmental conditions of the sample location, with the number at the tip corresponding to the individual sample. The names assigned by a taxonomic specialist and based on the geolocation data align with the tip labels. The major morphological characters used to help guide taxonomic identification are aligned with the tip labels, and the colors of the bar charts correspond to the sampling location.

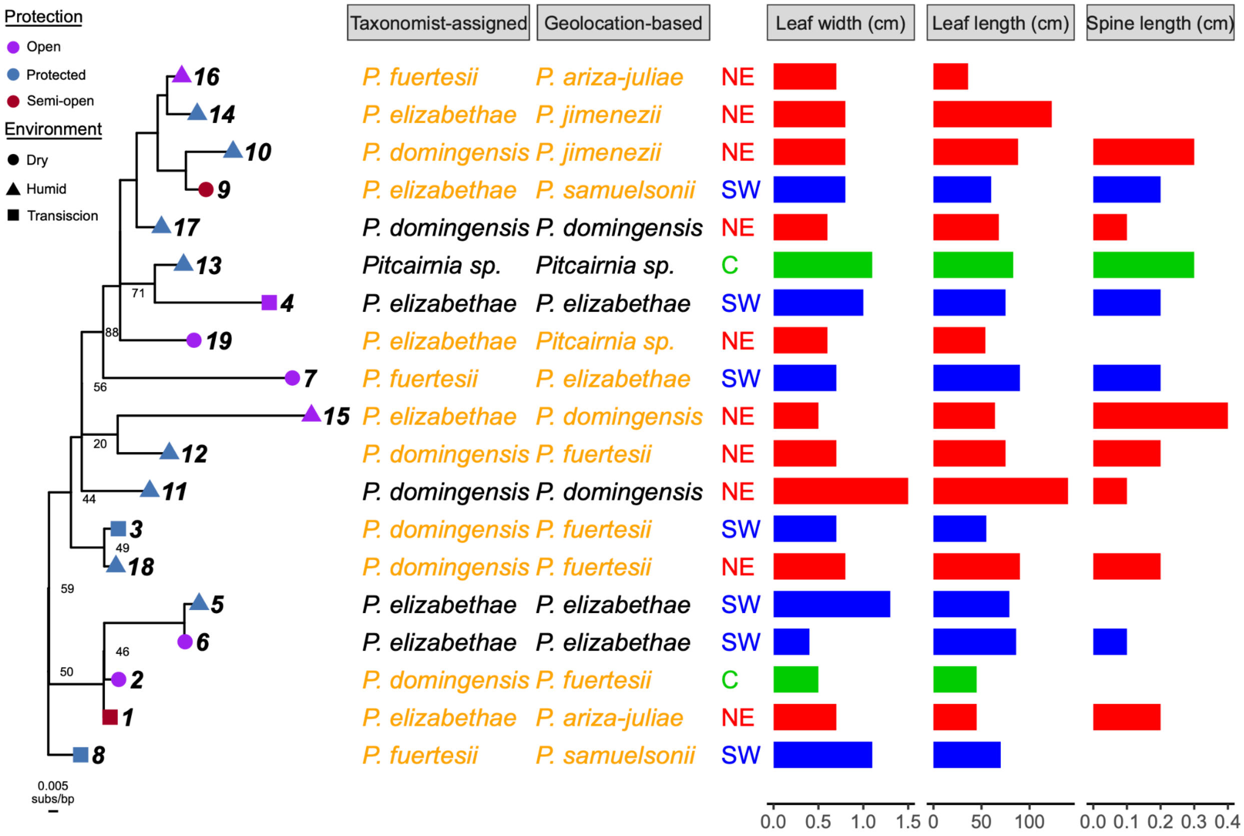

**Figure S3. The inferred relationships among individuals using Mitochondrion data.** The tree was inferred using an ML concatenated supermatrix, and support for the relationships is based on Ultrafast bootstraps, with the UFboot values for all poorly supported relationships (UFboot < 95%) labeled. The shapes and colors on the tips correspond to the environmental conditions of the sample location, with the number at the tip corresponding to the individual sample. The names assigned by a taxonomic specialist and based on the geolocation data align with the tip labels. The major morphological characters used to help guide taxonomic identification are aligned with the tip labels, and the colors of the bar charts correspond to the sampling location.

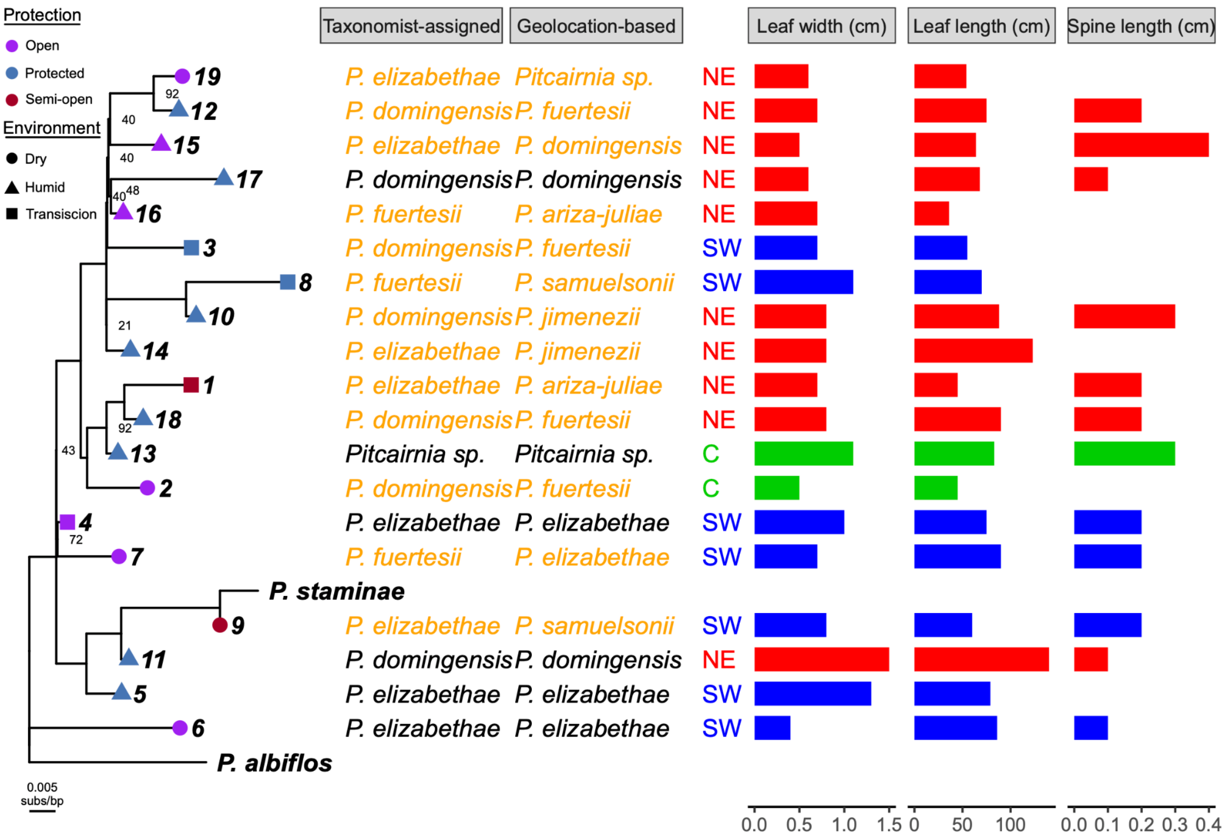

**Figure S4. Parsimony Informative Characters across the protein coding sequences for each dataset.**

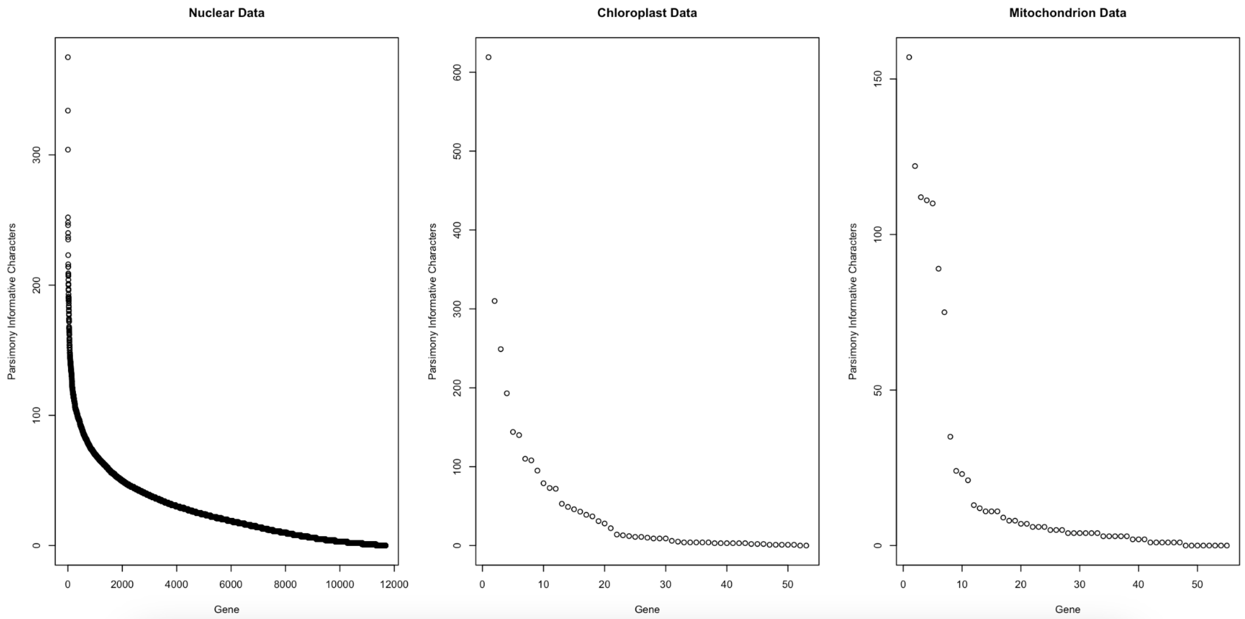

**Figure S5.** The distribution of allele frequency for the nuclear data compared to that as expected based on Hardy Weinberg Equilibrium.

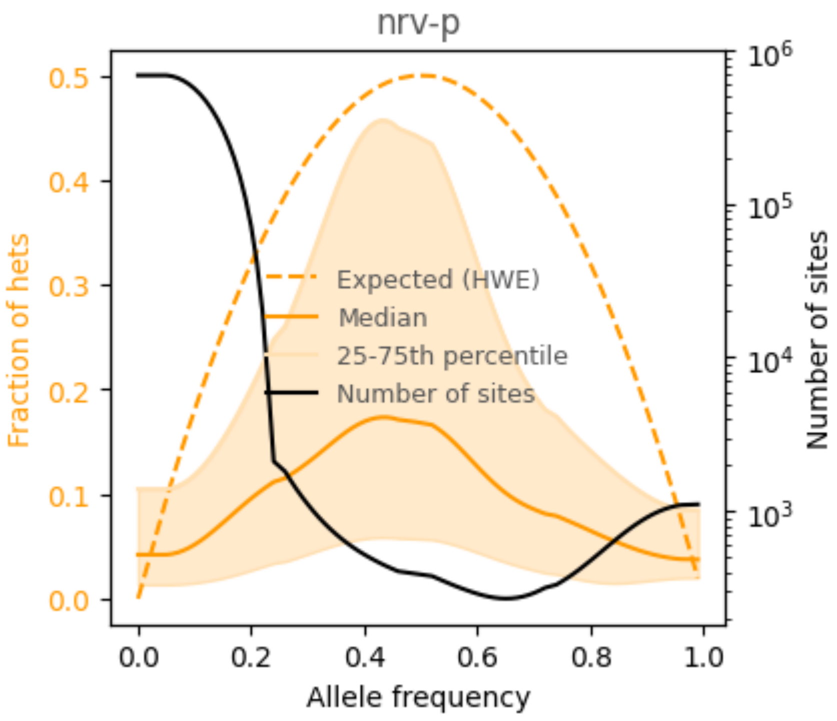

**Figure S6. Morphological differences among the named species of *Pitcairnia*.** Leaf length (top left), leaf width (bottom left), trichome density (top right), and marginal spine length (bottom right) for each sample ordered according to the specified metric. The ordered bars are colored according to taxonomist-assigned and geolocation-based naming schemes. Neither naming scheme shows a pattern with the measured morphological characteristics.

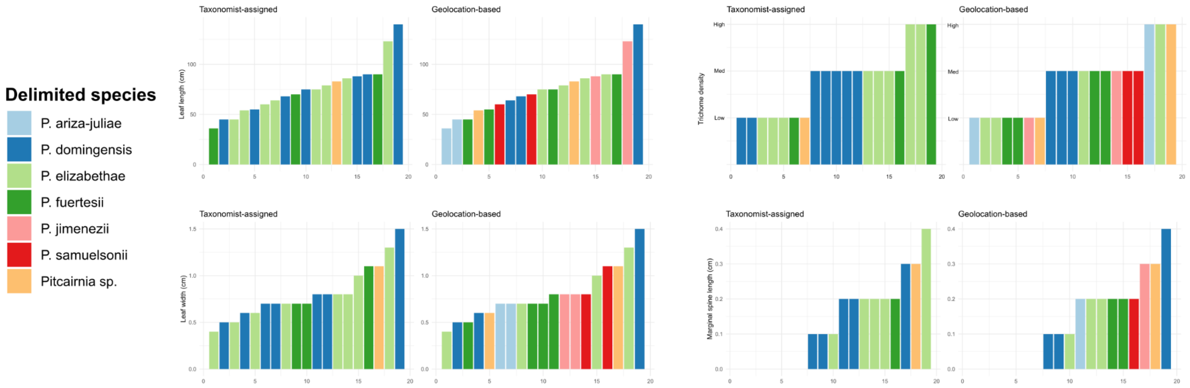

**Figure S7.** Examples of variation of two morphological characters measured: Size of marginal spines and trichome density found throughout the Dominican Republic.

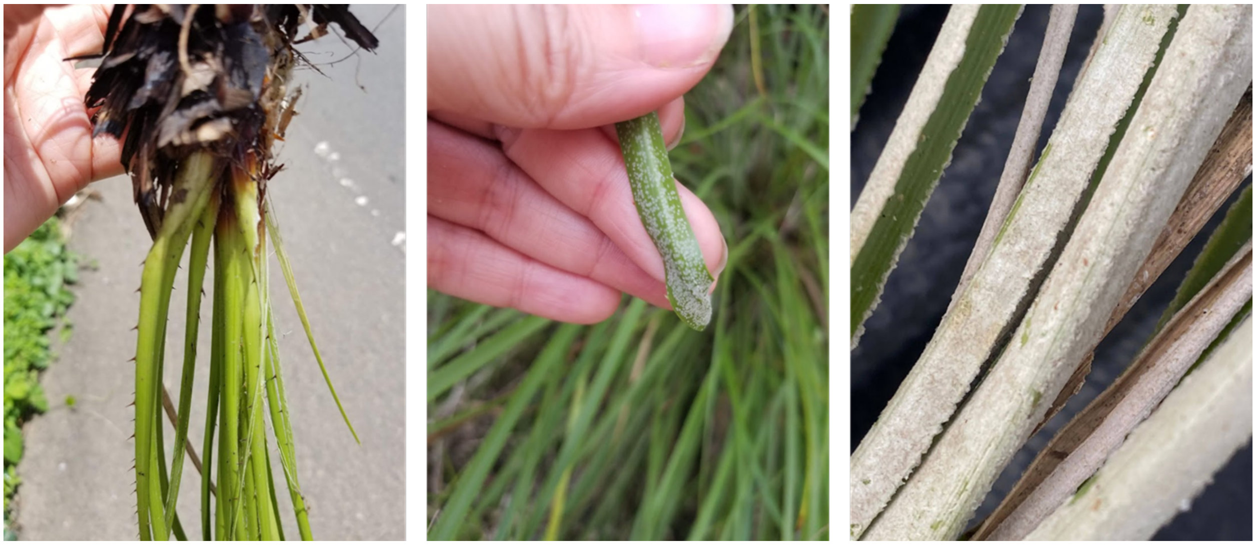

**Figure S8.** Although closely related, samples 11 (left) and 16 (right) show significant variation in size, trichome density, and leaf coarseness.

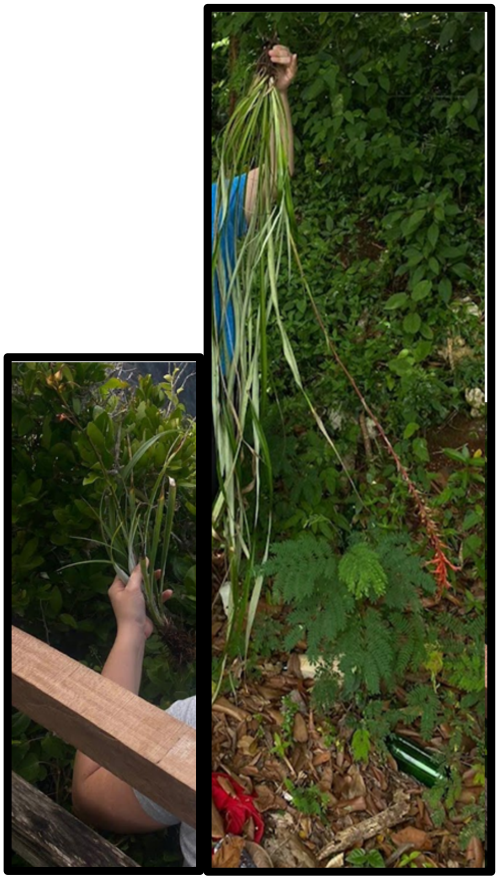

**Table S1.** Sample numbers, collections and their respective Herbarium number at National Herbarium of the Dominican Republic (JBSD).

| Sample No. | Collector | Herbarium No. |
|---|---|---|
| 01 | N. Ruiz-Vargas | RD23-01 |
| 02 | N. Ruiz-Vargas | RD23-03 |
| 03 | N. Ruiz-Vargas | RD23-04 |
| 04 | N. Ruiz-Vargas | RD23-05 |
| 05 | N. Ruiz-Vargas | RD23-09 |
| 06 | N. Ruiz-Vargas | RD23-11 |
| 07 | N. Ruiz-Vargas | RD23-14 |
| 08 | N. Ruiz-Vargas | RD23-17 |
| 09 | N. Ruiz-Vargas | RD23-18 |
| 10 | N. Ruiz-Vargas | RD23-19 |
| 11 | N. Ruiz-Vargas | RD23-22 |
| 12 | N. Ruiz-Vargas | RD23-01 |
| 13 | N. Ruiz-Vargas | RD23-04 |
| 14 | N. Ruiz-Vargas | RD23-18 |
| 15 | N. Ruiz-Vargas | RD23-16 |
| 16 | N. Ruiz-Vargas | RD23-12 |
| 17 | N. Ruiz-Vargas | RD23-21 |
| 18 | N. Ruiz-Vargas | RD23-10 |
| 19 | N. Ruiz-Vargas | RD23-06 |

**Table S2.** Full summary of sequenced samples and the resulting transcriptomes produced.

| Sample Name | Total RNA (ng) | RIN | Initial Reads | Nuclear Reads | Plastome Reads | Mitochondrion Reads | Nuclear trinity | Plastome trinity | Mitochondrion trinity | Nuclear transdecoder | Plastome transdecoder | Mitochondrion transdecoder | Nuclear final | Plastome final |
|---|---|---|---|---|---|---|---|---|---|---|---|---|---|---|
| 01 | 510 | 2.3 | 99484187 | 46578119 | 6349726 | 1682695 | 248894 | 5349 | 2146 | 36238 | 247 | 181 | 24990 | 38 |
| 02 | 840 | 4.3 | 77268317 | 46116739 | 865000 | 1407674 | 640144 | 887 | 3079 | 119112 | 163 | 240 | 23185 | 37 |
| 03 | 528 | 3.8 | 81670840 | 45084923 | 927284 | 1320711 | 1082079 | 1246 | 3804 | 194970 | 286 | 339 | 19864 | 36 |
| 04 | 4930 | 5.4 | 77788478 | 51905175 | 3207685 | 2675367 | 311360 | 4365 | 5143 | 50494 | 278 | 238 | 26490 | 47 |
| 05 | 510 | 3.2 | 89263036 | 49957936 | 5133436 | 1909637 | 314314 | 4753 | 2344 | 42470 | 351 | 178 | 25313 | 36 |
| 06 | 390 | 4.3 | 99040548 | 65979048 | 2596433 | 3218196 | 462755 | 4740 | 3346 | 17910 | 180 | 203 | 14762 | 35 |
| 07 | 2380 | 3.7 | 87455923 | 46613054 | 8306422 | 2209659 | 269078 | 10177 | 3194 | 41442 | 351 | 195 | 27264 | 49 |
| 08 | 1581 | 7.4 | 95356788 | 56232278 | 99537 | 2066630 | 564337 | 535 | 2862 | 142119 | 161 | 127 | 4500 | 39 |
| 09 | 6600 | 4.8 | 86434374 | 49579100 | 3901460 | 1678802 | 329667 | 3052 | 2533 | 42297 | 328 | 185 | 25067 | 44 |
| 10 | 3100 | 4.1 | 80217537 | 49375135 | 3585535 | 2127142 | 257662 | 2455 | 2147 | 40628 | 163 | 195 | 26311 | 42 |
| 11 | 1020 | 5.7 | 90185382 | 45720319 | 866775 | 1421561 | 389885 | 1461 | 2613 | 87365 | 276 | 262 | 20781 | 39 |
| 12 | 600 | 5 | 86422214 | 60871492 | 2105689 | 1678563 | 807519 | 2639 | 3165 | 180864 | 477 | 304 | 23180 | 38 |
| 13 | 1110 | 5 | 98358395 | 58213213 | 3565910 | 2246538 | 797692 | 3220 | 3359 | 142455 | 268 | 277 | 26284 | 41 |
| 14 | 899 | 4.9 | 95451297 | 52588180 | 1376271 | 1884013 | 652469 | 1413 | 3451 | 184269 | 171 | 326 | 24689 | 36 |
| 15 | 690 | 3.7 | 81388060 | 50583456 | 4039181 | 2913524 | 442691 | 4249 | 4416 | 62802 | 405 | 289 | 26586 | 47 |
| 16 | 990 | 4.1 | 84733604 | 49143522 | 4013399 | 2781999 | 387896 | 2580 | 2279 | 104925 | 288 | 281 | 24695 | 35 |
| 17 | 1488 | 5.7 | 85161150 | 45201696 | 478966 | 1656652 | 476894 | 472 | 3006 | 174964 | 98 | 300 | 18194 | 33 |
| 18 | 406 | 4.2 | 79558466 | 45963594 | 974380 | 2267451 | 714181 | 1886 | 3290 | 112723 | 497 | 286 | 23009 | 40 |
| 19 | 1290 | 6.1 | 88947285 | 52751861 | 1804642 | 1206667 | 1639473 | 2183 | 3016 | 340105 | 583 | 300 | 17837 | 38 |

| | | | | | | | | | | | | | | |
|---|---|---|---|---|---|---|---|---|---|---|---|---|---|---|
| *P. albiflos* | NA | NA | 56976403 | 40661866 | 325139 | 46110 | 97699 | 237 | 333 | 26558 | 99 | 73 | 19162 | 45 |
| *P. staminae* | NA | NA | 51790009 | 43995454 | 57717 | 9067 | 109488 | 235 | 166 | 30466 | 92 | 34 | 19752 | 9 |

**Table S3.** Plastome genes recovered.

| Gene | Number of transcriptomes with the gene |
|---|---|
| *accD* | 13 |
| *atpA* | 21 |
| *atpB* | 16 |
| *atpE* | 21 |
| *atpF* | 4 |
| *atpH* | 3 |
| *atpI* | 21 |
| *ccsA* | 13 |
| *cemA* | 6 |
| *clpP* | 13 |
| *matK* | 20 |
| *ndhA* | 21 |
| *ndhB* | 21 |
| *ndhC* | 9 |
| *ndhD* | 7 |
| *ndhE* | 6 |
| *ndhF* | 18 |
| *ndhG* | 3 |
| *ndhH* | 21 |
| *ndhI* | 4 |
| *ndhJ* | 11 |
| *ndhK* | 20 |
| *petA* | 7 |
| *petB* | 19 |
| *petD* | 11 |
| *petG* | 1 |
| *psaA* | 21 |
| *psaB* | 21 |
| *psaC* | 5 |
| *psaI* | 2 |
| *psbA* | 17 |
| *psbB* | 20 |
| *psbC* | 21 |
| *psbD* | 21 |
| *psbE* | 8 |
| *psbJ* | 1 |
| *psbK* | 16 |
| *psbM* | 1 |
| *psbN* | 2 |
| *psbZ* | 2 |
| *rbcL* | 16 |
| *rpl14* | 16 |
| *rpl16* | 16 |
| *rpl20* | 5 |
| *rpl22* | 19 |
| *rpl2* | 20 |
| *rpoA* | 13 |
| *rpoB* | 21 |
| *rpoC1* | 20 |
| *rpoC2* | 21 |
| *rps11* | 13 |
| *rps12* | 3 |
| *rps14* | 21 |

| | |
|---|---|
| *rps15* | 1 |
| *rps16* | 1 |
| *rps18* | 4 |
| *rps2* | 21 |
| *rps3* | 16 |
| *rps4* | 20 |
| *rps7* | 21 |
| *rps8* | 15 |
| *ycf1* | 21 |
| *ycf2* | 21 |
| *ycf3* | 2 |

**Table S4.** Mitochondrion protein coding sequences recovered.

| Protein Coding Sequence | Number of transcriptomes with the sequence |
|---|---|
| *cob* | 21 |
| *nad4* | 21 |
| *nad9* | 21 |
| *orf140-b* | 21 |
| *orf99-c* | 21 |
| *rpl2* | 21 |
| *rpl5* | 21 |
| *rps13* | 21 |
| *rps1* | 21 |
| *ccmC* | 20 |
| *CCMFC* | 20 |
| *CCMFN1* | 20 |
| *cox3* | 20 |
| *MATR* | 20 |
| *nad1* | 20 |
| *nad2* | 20 |
| *nad6* | 20 |
| *atp6* | 19 |
| *cox1* | 19 |
| *orf160* | 19 |
| *nad3* | 18 |
| *rps7* | 18 |
| *orf114-b* | 17 |
| *rps3* | 17 |
| *ccmB* | 16 |
| *nad4L* | 16 |
| *mttB* | 15 |
| *nad7* | 15 |
| *cox2* | 14 |
| *orf99-a1* | 13 |
| *orf111-b* | 12 |
| *atp4* | 11 |
| *ND5* | 11 |
| *orf186* | 11 |
| *atp1* | 10 |
| *atp8* | 10 |
| *orf173* | 10 |
| *orf25* | 10 |

| | |
|---|---|
| *orfB* | 10 |
| *ccmFN2* | 9 |
| *nad5* | 9 |
| *orf134-b* | 9 |
| *orf99-a2* | 8 |
| *orf127* | 7 |
| *atp1-a2* | 6 |
| *rps2A* | 6 |
| *rps2B* | 6 |
| *rps4* | 6 |
| *atp1-a1* | 5 |
| *orf158-a1* | 5 |
| *orfX* | 5 |
| *orf146-a* | 4 |
| *orf158-a2* | 4 |
| *rps12* | 4 |
| *rps2* | 4 |
| *atp9* | 3 |
| *orf101-a* | 2 |
| *orf117-c* | 2 |
| *orf105-a* | 1 |
| *orf110-c* | 1 |
| *orf140-a* | 1 |
| *orf99-b2* | 1 |
| *rps19* | 1 |
